# *Trypanosoma brucei* cattle infections contain cryptic transmission-adapted bloodstream forms at low parasitaemia

**DOI:** 10.1101/2025.02.21.639415

**Authors:** Stephen D. Larcombe, Edith Paxton, Christina Vrettou, Pieter C. Steketee, Keith R. Matthews, Liam J. Morrison, Emma M. Briggs

## Abstract

Tsetse-transmitted *Trypanosoma* parasites infect a wide host range and cause Human African Trypanosomiasis and Animal African Trypanosomosis in sub-Saharan Africa. The primary hosts of *Trypanosoma brucei* in tsetse fly endemic regions are non-human mammals, including agriculturally important cattle. In rodent infection models, *T. brucei* transitions from proliferative slender to tsetse-transmissible stumpy forms at high parasitaemia in a density-dependent quorum sensing-type process. However, chronic bovine infections are characterised by markedly lower blood parasitaemia levels; in most cases substantially below the density assumed to trigger slender-to-stumpy differentiation. This challenges the current (rodent-based) assumptions and quantitative parameter estimations around the generation of stumpy forms in the mammalian bloodstream by quorum sensing.

By combining scRNA-seq and microscopy in the first molecular characterisation of *T. brucei* forms in cattle blood, we observed mixed populations of parasites with slender and stumpy-like transcriptomes. The appearance of stumpy-like forms coincided with fewer proliferating parasites and parasites exhibited a shortened flagellum indicative of differentiation, despite the absence of an extreme stumpy morphology or developmental marker protein expression. Comparisons with slender and stumpy form transcriptomes from murine infection and *in vitro* culture demonstrated conserved transcriptomic signatures for both slender and stumpy-like forms in bovines, as well as host specific differences. These similarities and differences are key to understanding parasite development and transmission in its natural host.

## Introduction

Tsetse-transmitted African trypanosome parasites are a significant public health and socio-economic problem across 37 African countries where tsetse flies are endemic, due to the parasite’s ability to infect both humans and a wide range of domesticated and wild animals [1, 2]. The best studied trypanosome species, *Trypanosoma brucei*, is the causative agent of Human African Trypanosomiasis (HAT; subspecies *T. b gambiense* and *T. b. rhodesiense*), and one of the causes of African Animal Trypanosomosis (AAT or Nagana, *T. b. brucei*) in livestock. Thanks to extensive vector control efforts, human cases of trypanosomiasis are scarce relative to historic levels, but disease in cattle remains responsible for a substantial economic and humanitarian burden [3].

Most African trypanosome species depend on the tsetse fly vector to complete their life cycles, requiring these parasites to undergo a complex series of developmental changes when passing from a mammalian host to the insect vector [4]. Among African trypanosomes, *T. brucei* is unique in that its development to transmissible forms in its mammalian hosts is marked by a distinct morphological change; replicative ‘slender’ forms establish the infection, before differentiating into non-dividing ‘stumpy’ forms in a density-dependent manner [5, 6]. These stumpy forms are cell cycle arrested, relatively resistant to antibody-mediated complement lysis in comparison to slender forms [7] and exhibit molecular pre-adaptation for their onward development in tsetse flies [8]. Although slender forms can infect immunologically compromised and immature tsetse, stumpy forms preferentially infect adult tsetse flies, demonstrating their importance for disease transmission [9]. Indeed, parasites that have lost the capacity for stumpy formation (monomorphs) fail to complete development in the fly [10].

Most studies of slender and stumpy form dynamics in the mammal have employed rodent models of infection; here, parasitaemia levels in the blood undergo striking “peaks and troughs” in early infections, and parasitaemias can exceed 10^8^ parasites per ml, with infections often fatal within weeks [11]. Single cell transcriptomics (scRNA-seq) revealed that most parasites in these rodent infections, especially in the chronic phase when parasitaemia remains high, are either fully developed stumpy forms or differentiating intermediate forms [12]. Most intermediate forms are cell cycle arrested, express a stumpy-associated transcriptome profile, and are committed to stumpy-development, but are yet to develop the stumpy morphology [12]. Thus, morphology alone was not indicative of developmental status. Rodent models, however, are not reflective of cattle infections, or those of wild bovids and other species, which are characterised by low parasitaemia in the chronic phase of infection, which can persist for months to years especially under field conditions [13].This impedes detailed longitudinal study of *T. brucei* in cattle when parasitaemia falls below the detection threshold (usually by microscopy). Indeed, in classic studies using infections of different hosts, parasites were often only detectable following sub-inoculation in rodents, or through fly infection (e.g. [14]). Additionally, blood parasitaemia in chronic cattle infection persists well below the high density of parasites that is documented to produce morphologically stumpy forms in both rodent and *in vitro* models (>10^6^ parasites per ml). Consequently, how stumpy forms arise and support transmission at low parasitaemia remains a conundrum, and the parasite populations in bovine infections have not been characterised in detail or compared to those of murine or *in vitro* models – particularly in the context of chronic infection or with the extensive molecular knowledge of differentiation that has been generated in the last decade for *T. brucei*.

In this study, we conducted a molecular investigation of the development of *T. brucei* in the cattle bloodstream, with emphasis on parasite dynamics after the first waves of parasitaemia. We followed *T. b. brucei* infections for 60 days in calves and combined quantitative morphological parameters with scRNA-seq to characterise individual parasites within the potentially heterogeneous trypanosome population dynamically over time. We found that both slender and stumpy-like transcriptomic forms were present in varying proportions across infection, with stumpy-like forms reaching as much as 68% of the population in early infection. We also found few actively dividing parasites after day 23 of infection, consistent with a non-replicative population leading to low blood parasitaemia. Nonetheless, we found a paucity of parasites with classical stumpy morphology, or stumpy protein marker expression, despite their stumpy-like transcriptome. Instead, parasites have a morphology more closely resembling an intermediate/differentiating population. We also note host-specific differences between both slender and stumpy-like forms either generated *in vitro* or isolated from mice or cattle. By examining the developmental biology of trypanosomes in their natural hosts and in a phase of infection most representative of parasitaemias in the field, we provide new insight into how transmission to tsetse flies from cattle is supported.

## Results

### Parasitaemias in cattle comprise mixed populations expressing both slender- and stumpy-associated transcriptomes

Two calves were infected via the intravenous route with 1 x 10^6^ *T. b. brucei* Antat 1.1 90:13, and blood parasitaemia was monitored over 60 days (Figure 1A). Within the resolution of analysis, parasitaemia peaked between days 5 and 23 post infection, reaching an observed maximum of 4×10^4^/ml and 3×10^5^/ml in cows 1 and 2, respectively, before entering a chronic phase characterised by parasitaemia <1×10^4^/ml after day 23. In this chronic phase, for both cattle, parasitaemia persisted mostly between 1-3×10^3^/ml, or occasionally fell below the threshold of detection.

**Figure 1.**
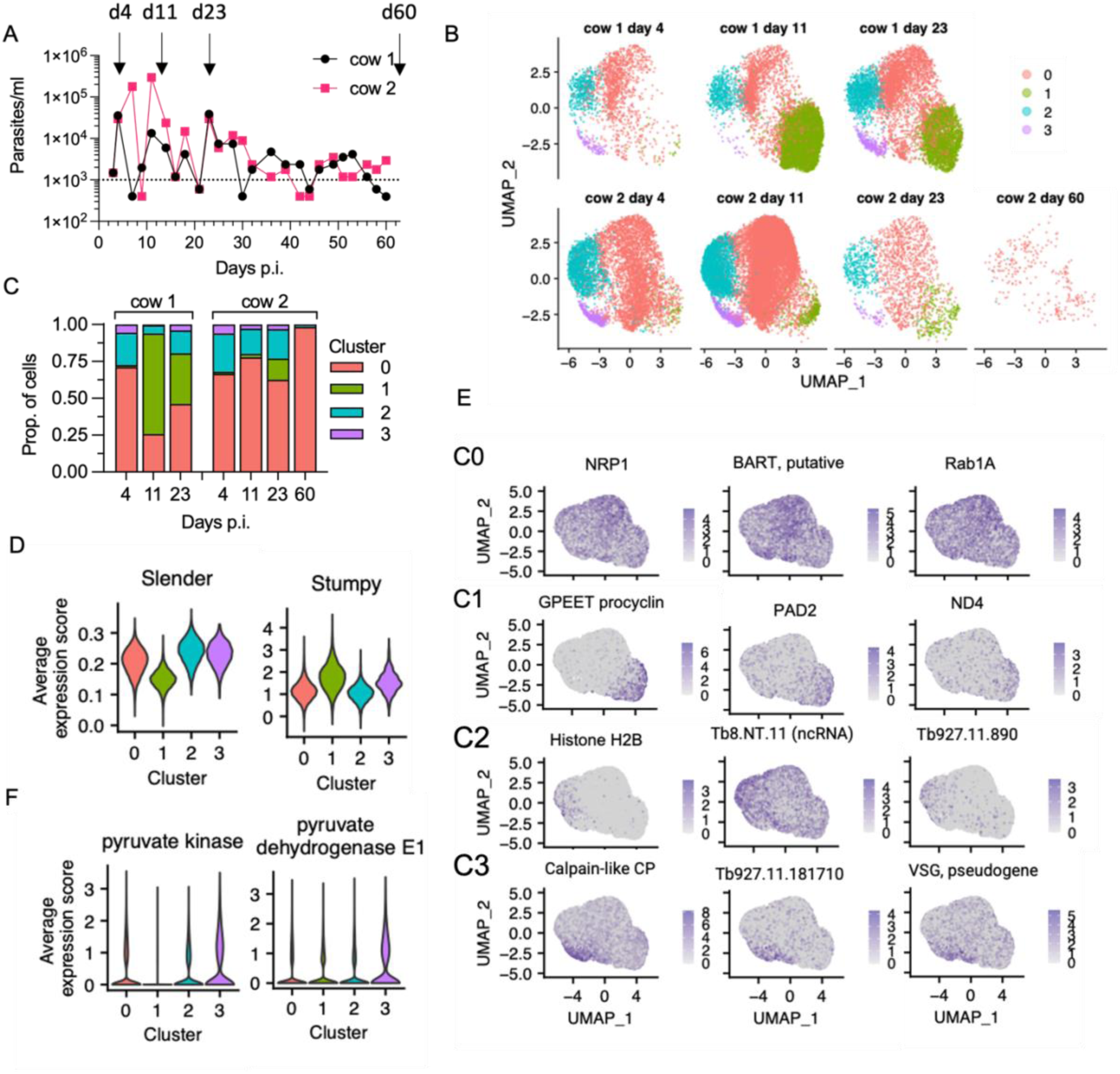
scRNA-seq identified slender- and stumpy-like forms of *T. brucei* in cattle blood. A) Blood parasitaemia levels for cow 1 (black) and cow 2 (pink). Dotted lines indicate low confidence threshold. B) Clusters of individual transcriptomes of *T. brucei* isolated on days 4, 11, 23 and 60 (only cow 2) post infection. Cluster colours are consistent across plots. C) Proportion of each sample, split by cow, in each of the clusters. D) Average expression scores of slender (left) and stumpy (right) marker genes for each cluster. Colours as in b. E) UMAPs coloured by expression level of three marker genes for each cluster. Gene IDs are shown for markers with unknown functions. F) Expression plots of two metabolic enzymes with highest expression in cluster C3

To investigate the potentially heterogeneous populations of *T. brucei* in the cattle bloodstream, we used scRNA-seq at time points when sufficient parasites could be isolated and purified; days 4, 11 and 23. Parasites were also purified from the blood at the end of the experiment (day 60) although extremely low parasite numbers at this time point hampered efforts to obtain sufficient cells for Chromium scRNA-seq analysis. Despite this, we obtained between 1,585 and 13,747 individual transcriptomes from *T. brucei* for both cows on days 4, 11, and 23, as well as 257 transcriptomes from cow 2 on day 60 (supplementary figure 2). Transcriptomes obtained from cow 1 on day 60 did not meet quality control thresholds for analysis and were excluded.

Samples were integrated together before dimension reduction and clustering analysis (Figure 1B). Four distinct clusters were identified and were detected in all samples, with the exception of cow 2 day 60 (Figure 1C) where the low number of transcriptomes captured limited analysis. Independent approaches identified that three of the clusters (C0, C2 and C3) had a transcriptome consistent with slender forms, whereas cluster C1 (green) contained transcriptomes that most closely resembled stumpy forms. Firstly, the average expression levels of previously identified slender-associated markers [15] was lower in cluster C1 compared to the other clusters and, conversely, the average expression of stumpy-associated markers was higher in cluster C1 compared to clusters C0, C2 and C3 (Figure 1D). Secondly, *de novo* identification of marker genes associated with each cluster via differential expression analysis followed by gene ontology (GO) term enrichment was used to identify biological processes associated with each cluster (supplementary figure 3, supplementary data 1). This revealed that Clusters C0, C2 and C3 had higher expression of genes associated with terms linked to proliferating slender forms: “glycolytic process” and “cytokinesis” [16]. In contrast, cluster C1 was associated with “translation” due to the high number of ribosomal protein transcripts that showed elevated levels in this cluster, as well as eukaryotic translation initiation factor 5A (EIF5A) and receptor for activated C kinase 1 (RACK1) [17] Notably, previous transcriptomic analysis of *T. b. rhodesiense* isolated at peak parasitaemia in rats, when stumpy forms are dominant, also showed high levels of translation associated transcripts [18]. Cluster C1 markers also included: Protein Associated with Differentiation (PAD) family members that are involved in the perception of the stumpy-to-procyclic differentiation signal [19]; procyclic form surface proteins EP and GPEET procyclin [20]; a mitochondrial NADH dehydrogenase subunit ND4; the stumpy-elevated and heat stress associated RNA regulator ZCH11 [21]; the iron responsive regulator RBP5 (Tb927.11.12100) [22]; and, the pteridine transporter Tb927.10.9080, which is a target of snoGRUMPY, a proposed regulator of stumpy formation [23]. Therefore, cluster C1 parasites expressed known transcript markers of stumpy forms, and were strongly associated with increased expression of genes required for global translation and developmental gene regulation.

Markers unique to each of the slender-like clusters C0, C2 and C3 (supplementary data 1) revealed heterogeneity in the slender forms identified in the cow bloodstream. Clusters C0 and C2 were defined by differences in their cell cycle phase signatures (supplementary figure 4), established using previously identified cell cycle marker transcripts [24]. Consistent with slender forms being proliferative, expression of S and G2/M phase marker genes was significantly increased in slender cluster C0 and even more so in slender cluster C2 (supplementary figure 4A), indicating that at least some parasites in the bloodstream were actively dividing at all of the timepoints we analysed. As enrichment for the G1 phase markers in any particular cluster was unclear (supplementary figure 4B), we could not confidently assign a threshold to estimate the proportions of parasites in each phase. However, stumpy-like cluster C1 had significantly lower expression of marker genes for each phase compared to the other clusters, most evidently for S and G2/M phase markers (supplementary figure 4A). Hence, cluster C1 parasites were most likely non-dividing and residing in G1/G0, consistent with their transcriptomic assignment as stumpy-like cells.

In addition to cell cycle phase signatures, cluster C3 was distinguished by a distinct set of marker genes relating to “tricarboxylic acid cycle” and “pyruvate metabolic process”. This subpopulation of slender-like forms had upregulated the expression of genes required to further metabolise pyruvate to generate ATP (Figure 1f), distinct from the classically described secretion of pyruvate by slender forms [25], but maintained expression of glycolytic components.

### The appearance of stumpy-like forms correlates with reduced proliferation and shorter flagella lengths

When possible, parasites were examined by microscopy, allowing the proportions of C1 parasites identified by scRNA-seq to be correlated with hallmarks of stumpy form development: cell cycle arrest, PAD1 surface protein expression and changed morphology (Figure 2).

**Figure 2.**
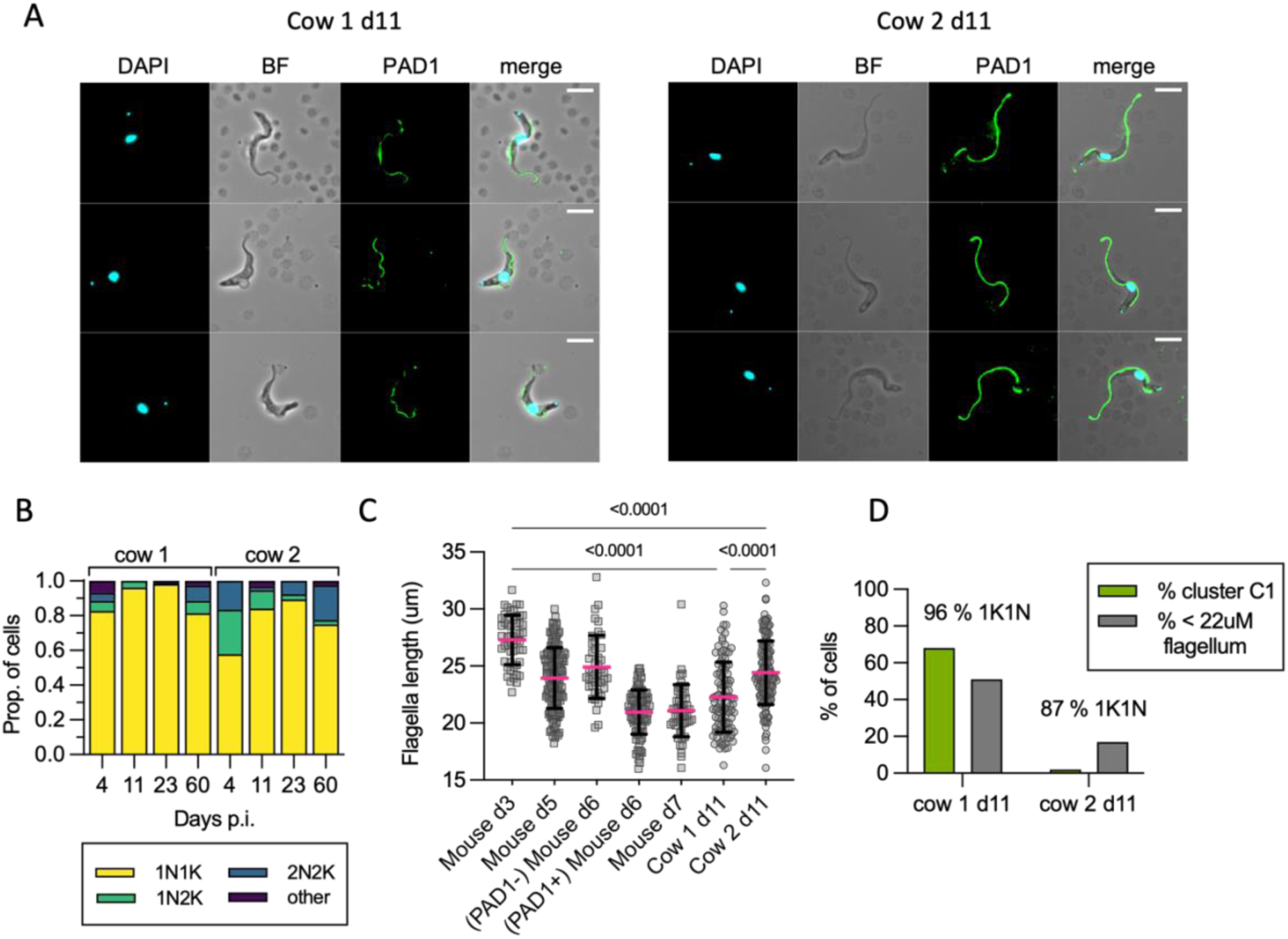
Cell Morphology and division status reflects scRNAseq cluster proportions. A). Representative cell images from parasites isolated on d11pi in cow 1 and cow 2, showing an absence of cell surface PAD1 staining (except the flagella). B) KN counts of parasites in each cow at each time point. C) The distribution of flagellar measurements from parasites in both cows on d11pi compared to controls taken from mice. On d3, all mouse parasites are slender, PAD1 negative, and fully replicative; on d5 all appear slender, are PAD1 negative, and only some are replicative, on d6 some parasites are PAD1 positive, very few are replicative; on d7 all are PAD1 positive and non-replicative. Parasites from the cows on d11 were significantly different from d3 true slender forms from mice, and from each other. P-values indicated show the result of Dunn’s multiple comparison tests between samples. D) A comparison of the proportion of cells belonging to stumpy-like cluster 1, and the proportion having a flagellar length most similar to mouse derived stumpy forms (<22um) showing the difference between cow 1 and cow 2. The differences are also reflected in the higher number of non-dividing (1K1N) cells in cow 1.

Dividing parasites first segregate the mitochondrial kDNA (K) followed by the nuclear genome (N) allowing cell cycle status to be inferred from the KN configuration. Parasites with segregated kDNA and 1 nucleus, 2K1N, represent those in late S phase or G2, whereas parasites with segregated kDNA and 2 nuclei, 2K2N, are post-mitotic. Proliferative parasites with 1 kinetoplast and 1 nucleus (1K1N) can be in G1 or early S phase, whereas stumpy cells have exited the cell cycle and uniformly exhibit 1K1N. The parasites isolated from cow 2 on d4, as the parasitaemia was rapidly ascending towards its peak in the first wave of infection, were enriched for dividing forms (43% 2K1N or 2K2N, Figure 2B). All other samples profiled contained 2K1N/2K2N parasites, albeit in low numbers, indicating that the cattle blood contained dividing *T. brucei* at each point but as a very small proportion of the overall population relative to the establishment phase. Notably, the proportion of 1K1N parasites reached 96% and 98% in cow 1 on days 11 and 23, respectively, indicating low levels of division consistent with stumpy development [12].These samples also contained the greatest proportions of cluster C1 parasites identified by scRNAseq analysis (Figure 1C), further evidencing that this population included abundant non-dividing stumpy-like forms.

Further attempts to quantify the proliferative proportion of parasites were made using an *ex vivo* plating assay, which previously allowed accurate detection of replication competent parasites when present as few as 0.1% of the total population [12]. Surprisingly, only 7% of parasites isolated from cow 2 on day 4 were able to replicate *ex vivo* despite the large proportion of actively dividing cells (as defined by 2K1N or 2K2N) and no other samples gave rise to proliferating cultures at all. Given the detection of active cell division markers by both microscopy and scRNA-seq in all cattle samples, it is reasonable to suggest that rapid adaptation to the bovine environment rendered the parasites unable to proliferate in culture media, regardless of their replicative capacity *in vivo*.

As the proportion of stumpy-like C1 parasites varied most between the two cows on day 11 (Figure 1C), we used this time point to correlate two established differentiation parameters: stumpy-specific protein PAD1 expression [19] and flagellar length[26]. No parasites showed expression of the PAD1 surface protein in either sample, but morphological differences between parasites in each were apparent. In mouse infections, flagellar length provides an indication of morphological differentiation, being shorter in stumpy forms, but there is also a quantifiable change between slender forms from proliferating populations (mean = 27µm) and differentiating populations (mean = 24 µm) that are yet to take on a full stumpy morphology (mean = 19 µm) [26]. Visual inspection of live parasites in the cattle revealed that most parasites had a slender to intermediate morphology (see supplementary video 3-1). Flagella length was used to quantify this morphological change and compared to the same parasite stabilate undergoing slender-to-stumpy morphological development in mice, as a means of calibrating this transition in a bovine infection with events occurring during the well characterised mouse model (Figure 2C). The shortening of the flagellum was evident during mouse infection, as expected, as the parasites transitioned from a proliferative slender population (day 3, mean length 27.3 µm) to a differentiating population of non-proliferating forms that do not yet resemble full stumpy forms (day 5, mean length 23.9 µm), a heterogeneous population of PAD1 negative differentiating forms (day 6, mean 24.9 µm) and PAD1 positive stumpy forms (day 6, mean 21 µm), before all cells are PAD1 positive stumpy cells on d7 (mean = 21 µm). Notably, most parasites isolated from both cattle on day 11 had a reduced flagellar length (means = 22.3 µm and 24.4 µm, respectively), more consistent with differentiating forms than proliferating slender forms. Additionally, the flagellum length distribution differed between cow 1 and cow 2 on day 11, with more parasites with shorter flagella in cow 1, consistent with a more differentiated population. In contrast in cow 2, with more proliferative parasites, flagella were longer. This correlates with the higher proportion of cells belonging to stumpy-like cluster C1 and with a 1K1N configuration in cow 1 than in cow 2 (Figure 2D).

Thus, the short flagellum, prevalence of cluster C1 cells, and those with a 1K1N profile, each are indicative of a prevalence of stumpy-like parasites in cattle blood that, at least in the samples investigated, did not express PAD1 protein, although the mRNAs of PAD genes were significantly enriched.

### Chronic infection parasite morphology is consistent with the presence of differentiating forms

Having established the cytological characteristics of the stumpy-like forms, we used these features to investigate the bloodstream forms present in chronic cattle infection when parasitaemia levels dropped to around 1×10^3^-1×10^4^ parasites/ml, below the level feasible for effective scRNA-seq analysis.

In this chronic stage, parasites were overwhelmingly 1K1N cells, particularly between days 49-58 (mean = 93.4% 1K1N, min = 89.4, max = 97.6) (Figure 3A). This was also observed when the parasitaemia was extremely low, indicating very low levels of active parasite division in the blood during most of the chronic infection. Despite the majority of parasites being non-dividing forms, only 2/824 parasites were observed that stained positively for PAD1, both from cow 2 on d58, which was consistent with the general absence of fully morphologically stumpy forms in the chronic phase (Figure 3B).

**Figure 3.**
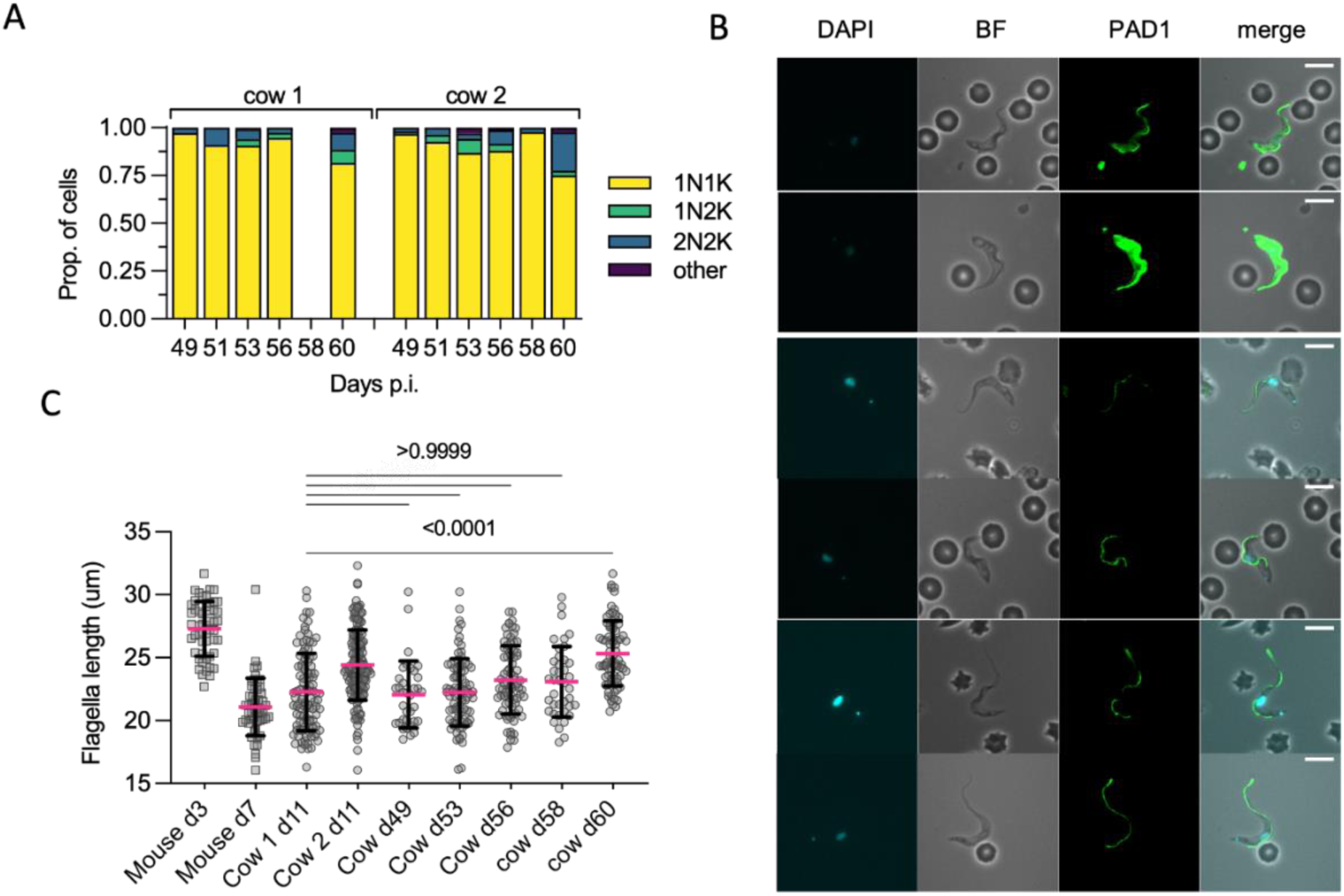
Chronic stage parasites have a morphology consistent with differentiation. A) Kinetoplast (K) and nucleus (N) configuration counts for each sample. B). Representative cell images from parasites isolated from d49-d60. Only two parasites (both shown) were scored as PAD1 positive, with most cells showing similar slender-intermediate morphology of those seen on d11. C) The distribution of flagella measurements in both cows in the chronic phase vs controls from d11pi and mouse slender forms (day 3) or stumpy forms (day 7). P-values indicated show the result of Dunn’s multiple comparisons tests between cow day 11 (which contains a high proportions of stumpy-like forms) and the samples taken in the late stage of infection.

To examine the developmental status further, the flagella length of chronic phase parasites was compared to that of cow 1 day 11, when a high proportion of stumpy-like parasites was evident (Figure 3C). No significant difference in flagella length was found between these time points, with the exception of day 60 when the mean flagella length was longer (mean for both cows =25.33 µm). Consistently, this time point was characterised by having a higher proportion of 2K2N cells in both cows, perhaps indicating a short replicative burst thereby enriching for slender forms at this time. Taken as a group, however, the flagellar length of chronic parasites was most similar to those from a non-dividing differentiating parasite population, supported by their similarity to the parasites from cow 1 on day 11 when the cluster C1 stumpy-like population was prevalent.

Thus, the chronic phase of infections was dominated by parasites with the shorter flagella associated with stumpy-like forms albeit without PAD1 protein expression, with a minority of actively dividing forms and scarce PAD1 positive morphologically stumpy forms.

### Host specific difference in *T. brucei* transcriptomes

To explore host-specific adaptations of the parasite populations, we compared the transcriptomes of parasites in each of the cow samples to our previous data derived from either *in vitro* culture or mouse infections by pseudobulking, whereby transcript counts for each gene were summed across every cell in the population within that sample. *In vitro* samples were mixed populations of slender and stumpy forms generated by exposure to brain heart infusion broth (BHI) to trigger stumpy development [15]. Mouse derived samples contained a mix of slender, differentiating and, most prominently, stumpy form parasites isolated on days 7 and 23 of infection [12].

For initial principal component analysis (Figure 4a), variant surface glycoproteins (VSGs) were removed from the genes used to calculate principal components to avoid grouping by VSG expression, which is expected to vary across infection. From the PCA analysis, it was apparent that *T. brucei* from mouse and culture systems were very similar to each other, although 55 genes have higher expression and 32 show lower expression in mouse samples when compared to parasites exposed to the BHI quorum sensing signal *in vitro* (Figure 4B, supplementary data 2). Of these, nine differentially expressed genes encoded different expression site associate genes (ESAGs) and VSGs, perhaps reflecting use of an alternative expression site. ESAG10, which encodes folate transporters, and ESAG6, transferrin receptor subunit, were overexpressed in mouse derived samples, whereas ESAG1 was downregulated compared to BHI generated samples.

**Figure 4.**
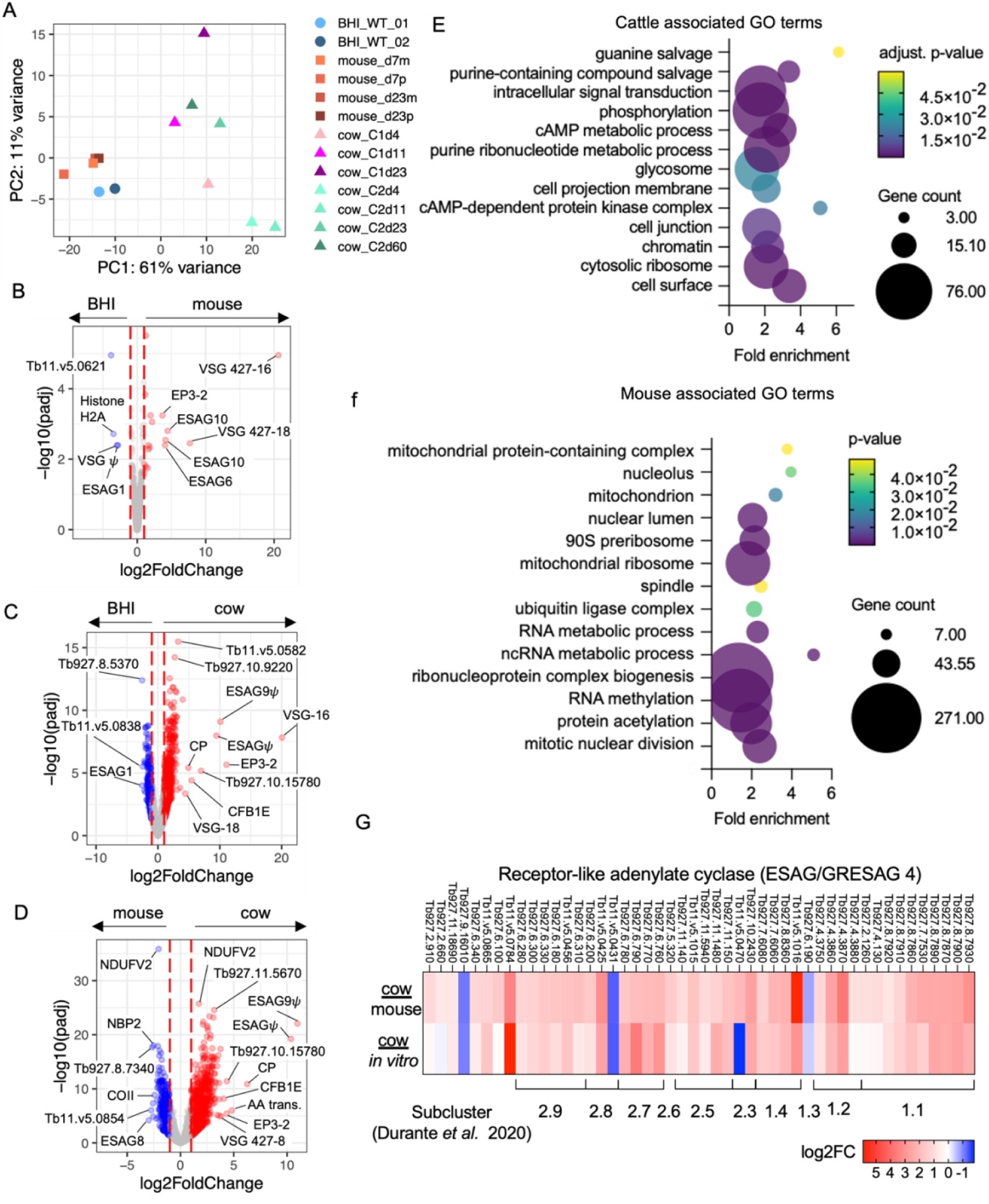
Host specific changes in *T. brucei* bloodstream forms. A) PCA plot of all *in vitro* (circles), mouse (squares) and cattle (triangles) derived samples. Volcano plots of adjusted p value (y-axis, - log10(padj) and fold change (x-axis, log2(fold change)) of differentially expressed genes between (B) BHI treated *in vitro* and mouse derived samples, (C), BHI treated *in vitro* and cattle derived samples and (D) mouse and cattle derived samples. E) Enriched GO terms in genes over expressed in cattle derived samples compared to mouse derived samples. F) Enriched GO terms in genes over expressed in mouse derived samples compared to cattle derived samples. G) Fold change (log2) in ESAG4/GRESAG4 transcript levels encoding receptor-like adenylate cyclase proteins, for parasites isolated from cattle relative to mice (top) *in vitro* culture (bottom). Clade and subcluster of each gene previously identified [27]is show below.

In contrast, the transcriptome of cattle derived parasites was clearly discriminated from those of mice and culture (Figure 4A). Most variance was explained by PC1 (61%), which separated cow samples from mouse or culture samples. Additionally, there was variation between samples taken over the course of cattle infection, and between individual cows at the same time point. Strikingly, after 23 days of mouse infection the transcriptome of cells remained closely related to cultured parasites, even when proportions of slender and stumpy forms varied between samples. In contrast, parasites from cattle could be discriminated from mice and culture derived parasites based on their transcriptome after only 4 days, demonstrating rapid gene expression changes, most likely related to adaptation, following bovine infection.

Cattle derived samples had far greater numbers of differentially expressed genes in comparison to both *in vitro* (Figure 4C) and mouse derived (Figure 4D) samples. GO term enrichment for genes with higher transcript levels in cattle derived samples (1,678 adjusted p < 0.05 and fold change >2), showed enrichment for transcripts associated with components of the cell surface, cytosolic ribosomes, chromatin, the flagellum and the glycosome (Figure 4E). Predicted cell surface proteins that varied in their expression between infections of mouse or cattle hosts included expression site associated genes (ESAGs). The ESAG classes that varied most commonly between host species were ESAG10 (3/7 annotated ESAG10 genes with detectable transcripts differentially expressed between mouse and cow), ESAG4 (7/18 genes), ESAG2 (9/25 genes) and ESAG8 (3/11 genes) (Supplementary data 2). The ESAGs with the greatest fold-change in transcript levels were two pseudogenes upregulated in cattle samples, ESAG9 (Tb927.11.110) and a non-classified ESAG (Tb929.11.130), that are located together in a likely VSG expression site (Figure 4D). The ESAGs with highest expression in mouse samples in comparison to cattle are also located the same VSG expression site, Tb427.BES112 (Figure 4D and supplementary data 2.) Strikingly, GRESAG4 (gene related to ESAG4), which are located in the core chromosomes and not the VSG expression sites, were also upregulated in cattle derived parasites (Figure 4G). The ESAG4 and GRESAG4 gene family encodes receptor-like adenylate cyclase proteins (ACs), which can be phylogenetically clustered into two clades and further subgroups [27]. Clades 1 and 2 contain ACs found to be translationally upregulated in bloodstream and procyclic forms, respectively, by ribosomal profiling [28]. Transcripts encoding ACs of both clades were upregulated in cattle derived trypanosomes (31.8% clade 1 and 33.7% clade 2) relative to samples from mouse infections and *in vitro* culture. Notably, transcripts encoding ACs termed ACP1-6 that are known to localise to the flagellum [29] and in some cases have been linked to social motility [30] were not differentially expressed between host environments. Thus, the observed changes in AC transcript expression levels do not appear to be linked to life cycle form development. An alternative possibility is that AC upregulation is linked to *T. brucei* adaptation or response to the host immune response. In early infection in mice, the activity of AC enzymes released by lysed parasites inhibits the production of trypanosome-supressing TNF-α in liver myeloid cells, allowing *T. brucei* to control host early innate immune [31].

### Host specific differences between developmental forms

Given the clear difference between hosts, we next sought to understand developmental form-specific differences between host environments. Attempts to fully integrate data from *in vitro*, mouse and cattle experiments were inconsistent across integration methods. Instead, a label transfer approach was used to predict the cell types in the cattle data (query dataset) using either the *in vitro* or mouse parasite cluster labels as the reference. Once the closest cell type match was found, markers for these clusters were compared. Providing evidence for the validity of using label transfer, a comparison of monomorphic slender form transcriptomes [24] to *in vitro* derived slender and stumpy forms [15] demonstrated that 97.9% of monomorphic slender forms labelled as slender, as expected (supplementary figure 5). Equally, and consistent with our earlier analysis (Figure 1d), predicting the cell types using the *in vitro* observed developmental forms as the reference resulted in the labelling of cattle cluster C1 as stumpy forms and clusters C0, C2 and C3 as slender forms (supplementary figure 6).

Previous analysis of *T. brucei* isolated from the mouse bloodstream on days 7 and 23 of infection [12] revealed that nearly all parasites were arrested stumpy-like forms at these time points, with only 2.1% of the captured parasites being slender forms (cluster M3, Figure 5A). In that study a clear stumpy population (cluster M0) was identified as the most common form in the mouse bloodstream (82.8%), and two other clusters (clusters M1 and M2) were identified that may represent “intermediate” forms transitioning between slender and stumpy extremes [12]. Using the label transfer method to predict the corresponding cell types between cattle and mouse derived parasite samples, we identified 70.9% of cattle cluster C1 (stumpy-like) as most similar to mouse cluster M0 stumpy forms (Figure 5B,5C). Smaller proportions of cattle cluster 1 were assigned as cell types M1 (7.3%) or M2 (5.3%), suggesting these putatively intermediate parasite forms were present but in low proportions in cows (Figure 3D). Therefore, the stumpy-like forms found in the cattle bloodstream were most similar in terms of their transcriptome to mature stumpy forms found in mouse infections, despite the differences in morphology and lack of PAD1 expression.

**Figure 5.**
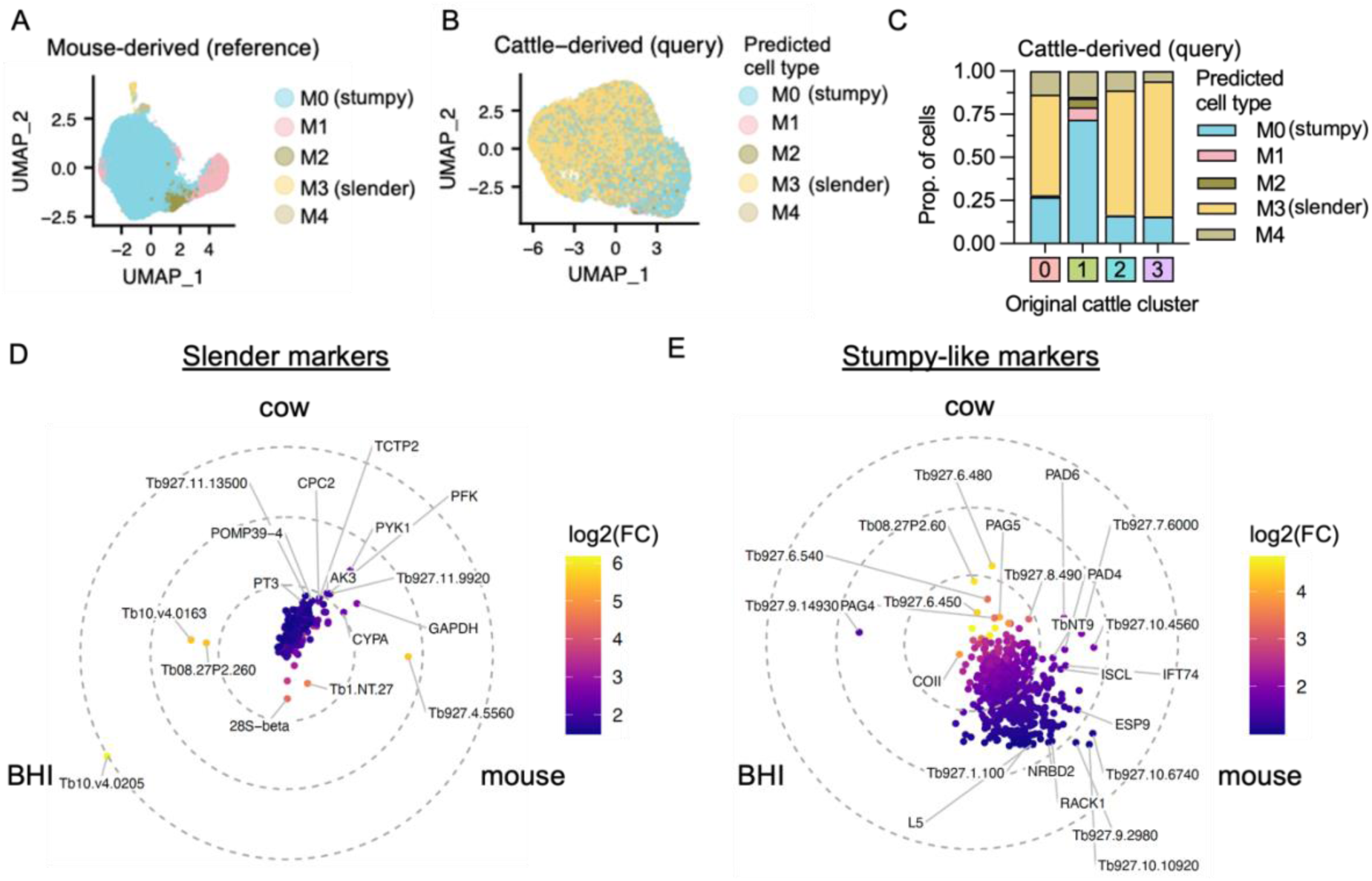
Comparison of *T. brucei* slender and stumpy-like isolated from mouse and cattle bloodstreams. A) UMAP of previously described scRNA-seq analysis of *T. brucei* isolated from the mouse bloodstream [12] used as a reference for label transfer. Slender (M3, yellow) and stumpy (M0, blue) clusters are highlighted. B). UMAP of cattle derived data coloured by predicted cell type. Labels and colours are consistent with reference data in Figure 2a. C) Proportions of each cattle derived cluster (x-axis) as shown in Figure 2b, predicted to be mouse derived cell-types. D) Similarity plot of the fold change in transcript levels of genes found to have higher expression in cow slender forms (clusters C0, C2 and C3 combined) and cow stumpy-like forms (cluster C1). Each point is one gene, colours by log2FC in transcript levels in cow data, positions by similarity across cow, mouse and *in vitro* BHI-generated data. Points to the centre show high similarity in fold change (slender over stumpy/stumpy-like forms) in every dataset, genes positions more distal to the have less consistent fold change between datasets. E) As in D, for cow stumpy-like form (cluster C1) markers relative to slender clusters. Fewer genes locate to the centre showing less consistency across datasets.

The transcript level fold change between stumpy/stumpy-like and slender forms was next quantitated for each dataset, and the similarity in these fold changes compared across the three conditions (Figure 5D, 5E, supplementary data 3). Genes located towards the centre of the plots had highly similar transcript fold changes across cow, mouse, and *in vitro* datasets, whereas those located closer to periphery showed varied fold changes between hosts. Slender marker genes across each dataset showed highly similar fold changes (Figure 5D), thus slender forms from cow, mouse and culture shared a common slender-associated transcriptomic profile. The top common markers consisted of genes linked to cell proliferation (e.g. histone H2B, chromosome passenger complex 1 and cytoskeleton components), numerous RNA binding proteins (e.g. RBP10, ZC3H38, ZC3H39, ZC3H31) and a non-coding RNA Tb1.NT.27.

The few genes that showed increased upregulation in slender forms from cattle and mouse infections compared to *in vitro* culture, included glycolysis enzymes (glyceraldehyde 3-phosphate dehydrogenase [GAPDH], pyruvate kinase [PYK1], ATP-dependent 6-phosphofructokinase [PRF]) indicating an increased emphasis on transcripts linked to glycolysis in both mammalian hosts compared to cultured parasites. This likely reflects the higher glucose levels in *in vitro* culture media (25 mM, [32]) compared to the mouse (7-9 mM, [33]) and cow (2-3 mM, [34]) bloodstreams. Other genes were VSGs, a putative polyubiquitin (Tb927.11.9920) and a paraflagellar rod component (Tb927.11.13500).

A common stumpy-associated transcriptome was also clear (Figure 5E), with the most conserved stumpy/stumpy-like markers including mitochondrial-encoded cytochrome oxidase subunit 2 (COII), procyclin proteins (EP1, EP2, GPEET, EP3-2), PAD2, multiple copies of retrotransposon hotspot protein 4 (RHS4) and several hypothetical proteins (see, supplementary data 3). Genes that showed higher transcript levels in stumpy-like forms in cattle, compared to stumpy forms isolated from mice or *in vitro*, included cAMP response protein 9 (CARP9; Tb927.8.4640), amino acid transporter 1 (AATP1; Tb927.8.7640), ESP9 (enriched in surface-labelled proteome protein 9; Tb927.9.11480), two putative metallopeptidase (FtsH33, Tb927.4.3300 and Tb927.11.3490) and adenylyl cyclase (GRESAG 4.4). Additionally, PAD4 (Tb927.7.5960), PAD6 (Tb927.7.5980) and PAD7 (Tb927.7.5990) were upregulated only in cattle derived stumpy-like forms.

## Discussion

Almost all of what is known about parasite development in mammalian trypanosome infections has come from *in vitro* culture and rodent models. Very little is known about how host and parasite interact to impact development in more clinically relevant systems. Here, we uncovered the presence of non-dividing stumpy-like forms that vary as a proportion of the cattle bloodstream population during infection, and persist into chronic infection at a significant level. Unlike rodent infections and *in vitro* models, the appearance of these forms was not linked to the highest parasitaemia levels, classical stumpy morphology or expression of the PAD1 marker protein. Yet, at almost all the timepoints we monitored, few parasites were visibly proliferative, and many had flagellar lengths consistent with the morphology of intermediate/differentiating forms. Thus, during a *T. brucei* infection the cattle bloodstream contains replication competent slender forms as well as a significant population of non-replicative stumpy-like parasites. Whether these latter forms are a) an intermediate stage committed to differentiation to classical stumpy forms, b) a reversible transient stumpy-like state that can revert to replicating slender forms, or c) a fully and terminally differentiated form that is distinct from morphological stumpy forms in mice, remains to be investigated. Nonetheless, these observations reveal several features of cattle infection dynamics, and pose new questions: firstly, why do parasites with a 1K1N configuration predominate at low parasitaemia despite the abundance of apparently replication-competent slender-like forms; secondly, if differentiation begins for a subpopulation of blood parasites, why are fully transitioned morphological stumpy forms rare; and, thirdly, do the stumpy-like forms we observe contribute to transmission?

The classic view of trypanosome infection dynamics entails undulating waves of infection where rapidly dividing slender forms expressing a dominant antigen type prevail, until either a density-dependent quorum sensing signal triggers differentiation into arrested stumpy forms or host immunity eliminates the bulk of the population, after which new antigen types repopulate the infection. This accurately reflects the first waves of parasitaemia, particularly in mouse infections, although a more complex view of VSG dynamics is now established based on higher resolution analyses of VSG expression profiles [35]. However, at parasitaemias below a density required to activate quorum sensing, only immune-mediated clearance of slender forms would be expected to operate in controlling parasitaemia. With the infection dynamics observed in cattle infections, according to parameters derived from rodent or *in vitro* systems, all parasites present should be replicative slender forms, which would maintain the infection through multiplication and the generation of antigenic diversity. However, our combined data suggest that the bloodstream population in cattle is heterogeneous despite the low parasitaemias; we find cell types that are configured in 1K1N (up to 97% in the chronic infection phase), have a shorter flagellum than would be expected for slender forms in the logarithmic growth phase [26], and express stumpy-associated transcripts. This invokes a requirement for the parasites to undergo developmental adaption in the mammalian host, even at low overall cell density in blood.

A number of scenarios could account for transmission-adaptation and arrest occurring in cattle, despite parasitaemia below the *in vitro* threshold for density-dependant differentiation (∼1×10^6/^ml). Firstly, it is possible that quorum-sensing pathways are triggered at lower density in cattle. In mice, *T. brucei* slender forms release oligopeptidase B and metallocarboxypeptidase 1 (among other peptidases) into the environment, generating the oligopeptide signal for stumpy formation [36, 37]. Although the level of transcripts for these peptidases was not seen to be elevated in our cattle samples, these enzymes or their oligopeptide products may be more active, or more stable in cattle, allowing development at lower parasitaemia. Secondly, stumpy-like form development may be triggered by alternative signals that are not only dependent upon parasite density. For example, an inflammatory environment may generate an oligopeptide signal driving development [38], or an environmental metabolite may promote stumpy formation in a quorum sensing-independent mechanism. Thirdly, parasites may achieve much higher density outside the blood in cattle, with stumpy-like parasites then seeding into the circulation. This latter scenario in particular has some experimental support, as analysis of mouse infections has previously demonstrated that there are apparently insufficient proliferative bloodstream forms to sustain the parasitaemia, with proliferation and development predicted to be occurring in the tissues [12]. This is further supported by the appearance of novel VSG types in blood after their emergence in the extravascular spaces [39]. The tissue-distribution of *T. brucei* in cattle is poorly understood [40] but recent diagnostic work on tissues from cows in Ghanaian abattoirs indicates that more positives are found in tissues than in blood alone [41].

The absence of fully morphologically stumpy forms in the cattle blood may be contributed to by an inability to successfully enter the circulatory system from the tissues, or by a more limited progression through the hierarchy of events that generate classic stumpy forms. In earlier analyses, we mapped the events of stumpy formation *in vivo* in mice, revealing that a stumpy transcriptome precedes cell cycle arrest, PAD1 expression and morphological development to stumpy forms [12]. Similarly, in cattle, parasites may progress along the pathway to stumpy formation but not fully develop to its latter stages, potentially predominating as uncommitted forms with the developmental flexibility to return to proliferation as slender forms or commit to arrest as stumpy-like forms. Entry into the fly may then activate PAD protein expression, which is thermoregulated in the case of PAD2 [19], to initiate development to procyclic forms from cluster C1 parasites with elevated PAD mRNA. Unfortunately, the inability to culture parasites *ex vivo* after cattle infection has limited the ability to explore whether parasites are committed to arrest or can return to proliferation; however, the molecular basis of this adaptation will be an interesting area of exploration. An alternative explanation for the lack of morphologically stumpy forms in cows may be differences in parasite clearance rates between hosts. Immune exhaustion caused by loss of B cells has been demonstrated in chronically infected mice, leading to reduced ability to remove parasites and their eventual death [42], this occurring at a time at which mice appear to have reduced ability to clear specific antigenic variants [35]. If parasites can survive longer before their removal in mice than cows, then a full morphological transition, which may take days to complete, would be more likely to be observed in mice.

Our results highlight differences between mouse and cattle infections, and between early and chronic phases. In the early stage of infection, mouse and cattle infections both show relatively high initial parasitaemia, which is controlled by the host. However, subsequently, cattle infections differ in being at much lower parasitaemia and show less evidence of clearly discriminated slender and stumpy morphotypes. These distinct acute and chronic phases may serve to promote establishment through the rapid proliferation of slender forms early on, where higher antigenic diversity serves to overcome any pre-existing field immunity and parasite growth can be restricted by quorum sensing in the first peak. Subsequently, the chronic phase would be comprised of parasites that can either sustain infection through proliferation and antigenic diversity or be adapted for transmission [43]. These “pre-committed” cell types that predominate in chronic infections might reflect a bet-hedging strategy for the parasites, optimised for survival or transmission where tsetse uptake is uncertain and long-term infection maintenance is required.

The distinctions between parasite transcriptomes in mice and cows were also evident as early as day 4 post infection, highlighting rapid gene expression changes, most likely related to adaptation to the host environment. This was also supported by our inability to culture parasites *ex vivo* from cows beyond this time point, despite clear evidence that large proportions of these parasites had a slender-associated transcriptome consistent with replicative capacity. The gene expression profiles of culture and mouse derived cells were in contrast much more similar, and it is notable that mouse-derived *ex vivo* cells will readily grow in the conditions attempted with cattle derived cells. The few genes with differing transcript levels in cattle slender forms compared to mouse and culture derived slender forms were largely linked to glycolysis, perhaps suggesting a switch in energy metabolism dynamics in response to the highly diverging host environments. However, such rapid adaptions will likely also involve protein level changes that could not be investigated in this study.

In summary, we have for the first time characterised the parasite populations that comprise acute and chronic phases of trypanosome infections in cattle and related these to established paradigms in mice and *in vitro*. We observed that infections are sustained at low parasitaemia and comprised of parasites that are not morphologically stumpy but express transcripts indicative of that developmental form. By establishing the molecular characteristics of parasites from the chronic phase of the infection, we have also extended beyond previous rodent-based laboratory studies of acute infections and broadened analysis to parasites in chronic infections in the natural host, representative of most infections in the field.

## Materials and methods

### Ethics statement

Animal experiments were carried out in the Large Animal Research and Imaging Facility at the Roslin Institute, University of Edinburgh, under United Kingdom Home Office Project License number PE854F3FC. Studies were approved by the Roslin Institute (University of Edinburgh) Animal Welfare and Ethical Review Board (study numbers L447 and L475). Care and maintenance of animals complied with University regulations and the Animals (Scientific Procedures) Act (1986; revised 2013).

### Cattle infections, parasitaemia and microscopy

Two male post-weaning Holstein Friesian calves aged 4-6 months were housed in vector proof containment (BSL2) conditions at the Large Animal Research and Imaging Facility of the Roslin Institute, University of Edinburgh. To initiate calve infections, blood containing *T. b. brucei* Antat 1.1 90:13 was freshly harvested from an infected donor mouse and quantified. Blood containing 1×10^6^ parasites in 1 ml from this same stock was used to infect each calf via the intravenous route. As blood parasitaemia was frequently extremely low and below the limit of accurate quantification using buffy coat preparation, we used a conversion between buffy coat and haemocytometer counts (parasites/ml = 29478 x parasites/field of view [FOV]) on all data points to infer parasitaemia at time points where parasites were detected but in extremely low numbers (<1 parasite per buffy coat FOV; Supplementary Figure 1).

### Parasite purification and scRNA-seq

200 ml of blood was collected for each scRNA-seq sample on days 4, 11 and 23. To obtain a pure parasite sample, a combination of red blood cell (RBC) lysis and DE52 column purification was used. Each whole blood sample was aliquoted into eight 50 ml falcons (∼16 ml per falcon) and centrifuged at 1,200 x *g* for 15 min without breaks to separate blood. The plasma was carefully removed, leaving RBCs and buffy coat behind where parasites will be located. A hypotonic lysis buffer (0.3 % NaCl in H_2_O) was added to each tube to a total volume of 45 ml and inverted. 5 ml of 10X PBS (Phosphate Buffered Saline) was then added and tubes inverted to halt RBC lysis. Samples were centrifuged at 80 x *g* for 20 min to pellet trypanosomes and retain as much of the blood cells in the supernatant as possible. Supernatant was then removed, and each pellet was combined by resuspending all in 10 ml of PSG (1X PBS + 1% D-glucose) and combining into one 50 ml tube. The combined sample was centrifuged at 400 x *g* for 10 min and supernatant removed, leaving 2 – 3ml of volume to resuspend the pellet in. The resuspension was mixed 1:1 with DE52 cellulose slurry and then applied to a glass filter column containing ∼100 ml of DE52 cellulose pre-washed with warm PSG. Columns were slowly washed with ∼100 ml of PSG to obtain pure trypanosomes and retain blood cells in the columns. Eluted parasites were then centrifuged at 400 x *g* for 10 min to pellet. Supernatant was removed to leave ∼1ml of PSG to resuspend the pellet. Parasites were then counted with a haemocytometer and adjusted to 1000 cell/µl by transfer to a clean Eppendorf, further centrifugation (400 x *g* for 10 min) and resuspension in an appropriate volume of PSG. Cells were counted again and adjusted as needed before 20,000 were loaded onto the Chromium (10X Genomics) controller for scRNA-seq. Sequencing was performed with Illumina NextSeq 2000 to generate 28 bp x 130 bp paired reads to a depth of at least 63,415 mean reads per cell. In the case of cow 2 day 23, two libraries were repaired and sequenced as a low droplet formation was apparent in this sample.

### scRNA-seq processing and cluster analysis

scRNA-seq data were mapped to the *T. brucei* TREU927 reference genome with extended UTRs to ensure all reads mapped were correctly assigned to a gene coding region. The kDNA maxicircle and *T. brucei* Lister 927 telomeric expression site contigs were also included to create a “hybrid” reference, as these regions are not present in the TREU927 reference [44]. Mapping was performed with Cellranger count and default settings to generate the genes counts matrices. Each sample was inspected and filtered to remove outliers that are likely poor-quality transcriptomes or multiplets (Supplementary Figure 2). Outliers with high kDNA (mitochondrial) and ribosomal RNA transcripts were also removed as these both indicate poor quality transcriptomes.

Each sample was normalised, and log transformed with Scran[45] independently. Variable genes were identified by both Scran and Seurat[46] methods and the common genes (with VSGs removed) were assigned as variable genes for each sample. Cattle samples were integrated using Seurat, scaled and PC analysis performed. UMAP was performed for visualisation using the top 20 PCs, as was nearest neighbours and clustering analysis.

For slender, stumpy and cell cycle phase scoring, average expression score of previously identified marker genes lists [15, 24] was calculated using the Seurat function *AddModuleScore*. For assigning the most likely cell cycle phase with these markers, scores were scaled between 0 and 1 was comparison and the phase with the highest expression score was assigned to each transcriptome. If all phase scores failed to reach the threshold of 0.5, the transcriptome was “Unlabelled”.

Cluster marker genes were identified using MAST[47] differential expression tests to find genes over expressed in one cluster compared to all other clusters. GO term analysis was performed via the TriTrypDB[48] website.

### Cross scRNA-seq dataset comparisons

The scArches package[49] was used to identify the cell types in cattle derived transcriptomes based on *in vitro* and mouse derived slender and stumpy *T. brucei* cell types that had characterised previously. Independently, reference datasets were normalised and logp1 transformed with Scanpy[50]. 1,500 variable genes were selected with Scanpy and the data subsetted to only use counts from these genes. A model was then created with scPoli [51] and trained on the reference data set (*in vitro* or mouse derived) using previously defined clusters as the cell type key. Models were then used to predict cell type in the cattle query data. To calculate fold change in transcript levels for all genes between slender and stumpy/stumpy-like clusters, FindMarker() was used with all filters set to 0, and the test.use set to “MAST”. Top markers for each cluster were defined as having an adjusted P value < 0.05, an average log2(fold-change) > 0.5 and being detected in at least 5% of the cells in the cluster under investigation and was limited to the top 500 after VSGs were removed.

### Pseudobulk RNA-seq analysis

To directly compare cattle data to previous scRNA-seq datasets of *in vitro* and mouse derived slender and stumpy forms, these were remapped to the “hybrid” reference genome and pre-processed in the same steps used in previous publications (where the telomeric contigs were not included for mapping). For consistence all QC filtering was performed exactly as previously descript for each. For each sample, total counts for each gene across all cells in the sample were summed to generate one total transcript count per gene. VSGs and contaminating ribosomal RNA transcripts were removed before DESeq2[52] was used for PC analysis using the top 2,000 genes. Individual DESeq2 differential expression tests were performed to compare conditions.

## Data availability

All scRNA-seq sequences is deposited on the European Nucleotide Archive under study accession number PRJEB66078.

## Code availability

All code needed to reproduce this analysis is accessible at zenodo.org under DOI:10.5281/zenodo.14515536.

## Acknowledgements

This work was supported by a Wellcome Trust Investigator award to KM (221717/Z/20/Z), a Wellcome Trust collaborative award to KM and LM (206815/Z/17/Z), a Sir Henry Wellcome Fellowship awarded to EB (218648/Z/19/Z) and a Roslin Institute Pump Priming BBSRC grant to LM, PS, KM and EB (sub-award from BBS/E/RL/230002C). LM, EP, CV and PS are funded through core support to the Roslin Institute by the United Kingdom Biotechnology and Biological Sciences Research Council (BBS/E/RL/230002C); PS is supported by a BBSRC Discovery Fellowship (BB/X009807/1); EB is supported by an MRC Career Development Award (MR/Z504786/1). We would like to acknowledge the assistance of staff members at the Large Animal Research and Imaging Facility, University of Edinburgh; specifically, James Nixon, Peter Tennant and Adrian Ritchie, who provided invaluable expertise and assistance in the experimental cattle infections. We would also like to thank Stefano Guido of the University of Edinburgh for expert advice, and Richard McCulloch and his group at the University of Glasgow for accommodating the single cell experiments. Finally, we would like to thank Julie Galbraith of Glasgow Polyomics for facilitating this unpredictable study and providing expert scRNA-seq library preparations and sequencing.

## Author contributions

PS, EB and SL developed the approach for parasite isolation. EP, CV, EB and SL produced the data in this study. SL performed microscopy analysis; EB performed scRNA-seq analysis. SL and EB produced the manuscript, along with KM and LM and comments from co-authors. All authors contributed to the interpretation of results, discussion and final manuscript.

## Competing Interests statement

The authors declare no competing interests.

---

**Supplementary figure 1.**
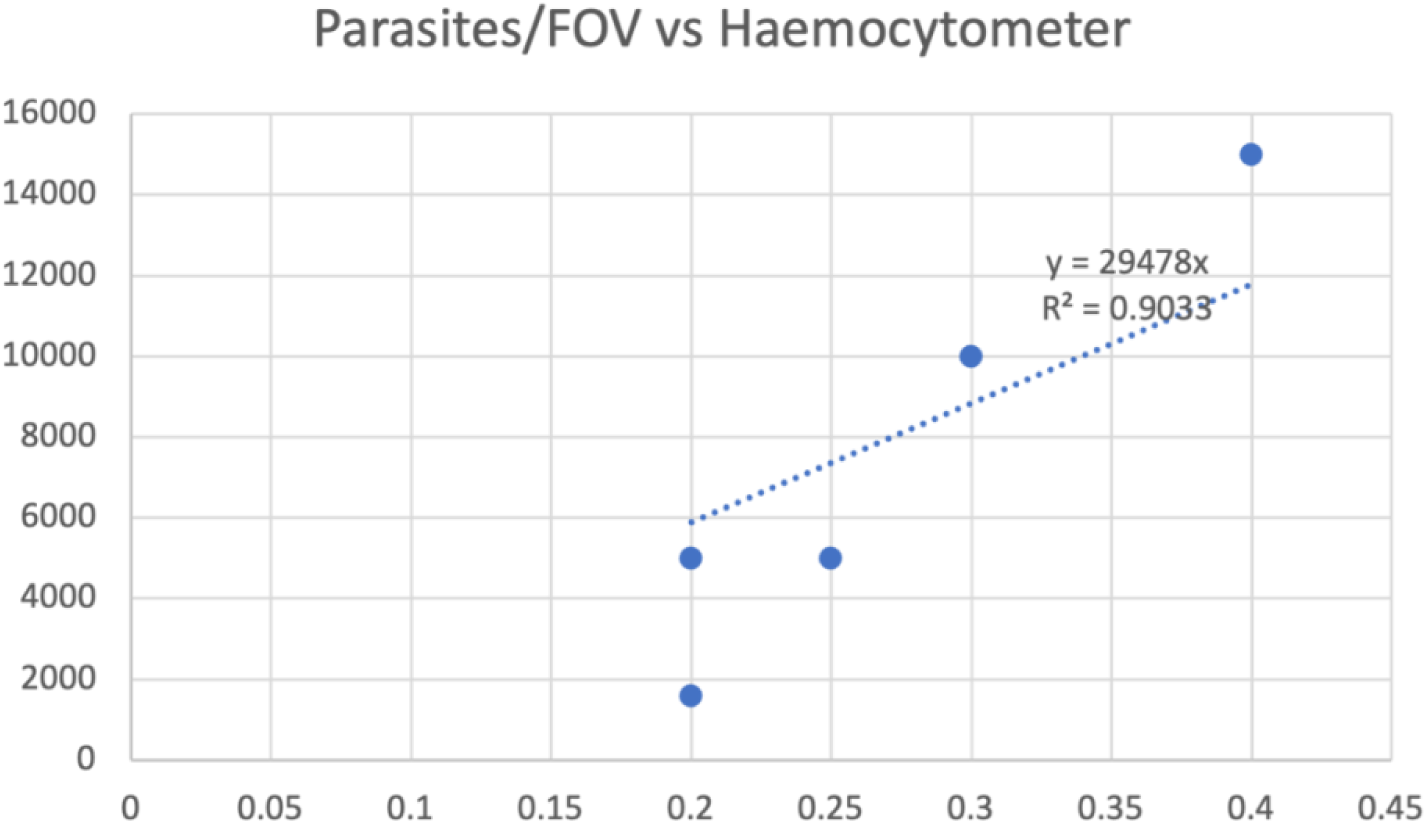
Number of Buffy Coat Parasites/FOV vs Parasites/ml based on counts where fewer than 1 cell / field of buffy coat was observed. R2 = 0.9 Equation: Parasitaemia/ml = 29478xFOV.

**Supplementary figure 2.**
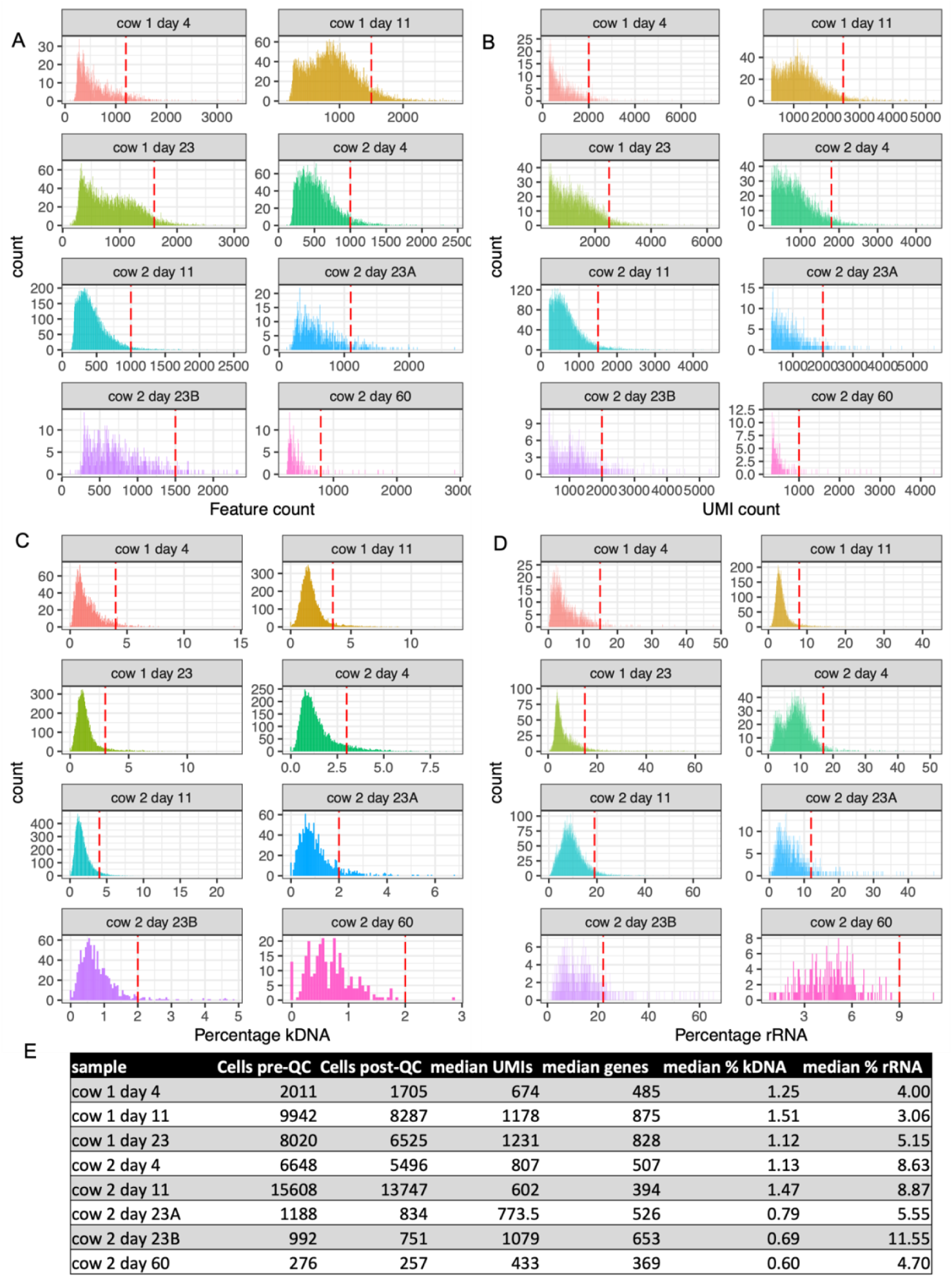
Metadata associated with scRNA-seq samples. Histograms showing the number of genes for which transcripts are detected (features, A), unique transcripts (UMIs, B), percentage of transcripts expressed from the mitochondrial kinetoplast maxicircle genome (Percentage kDNA, C) and percentage of transcripts encoding ribosomal RNA (Percentage rRNA, D), per parasite transcriptome. Red dashed lines indicate quality control thresholds used. E) Summary table of QC measures

**Supplementary figure 3.**
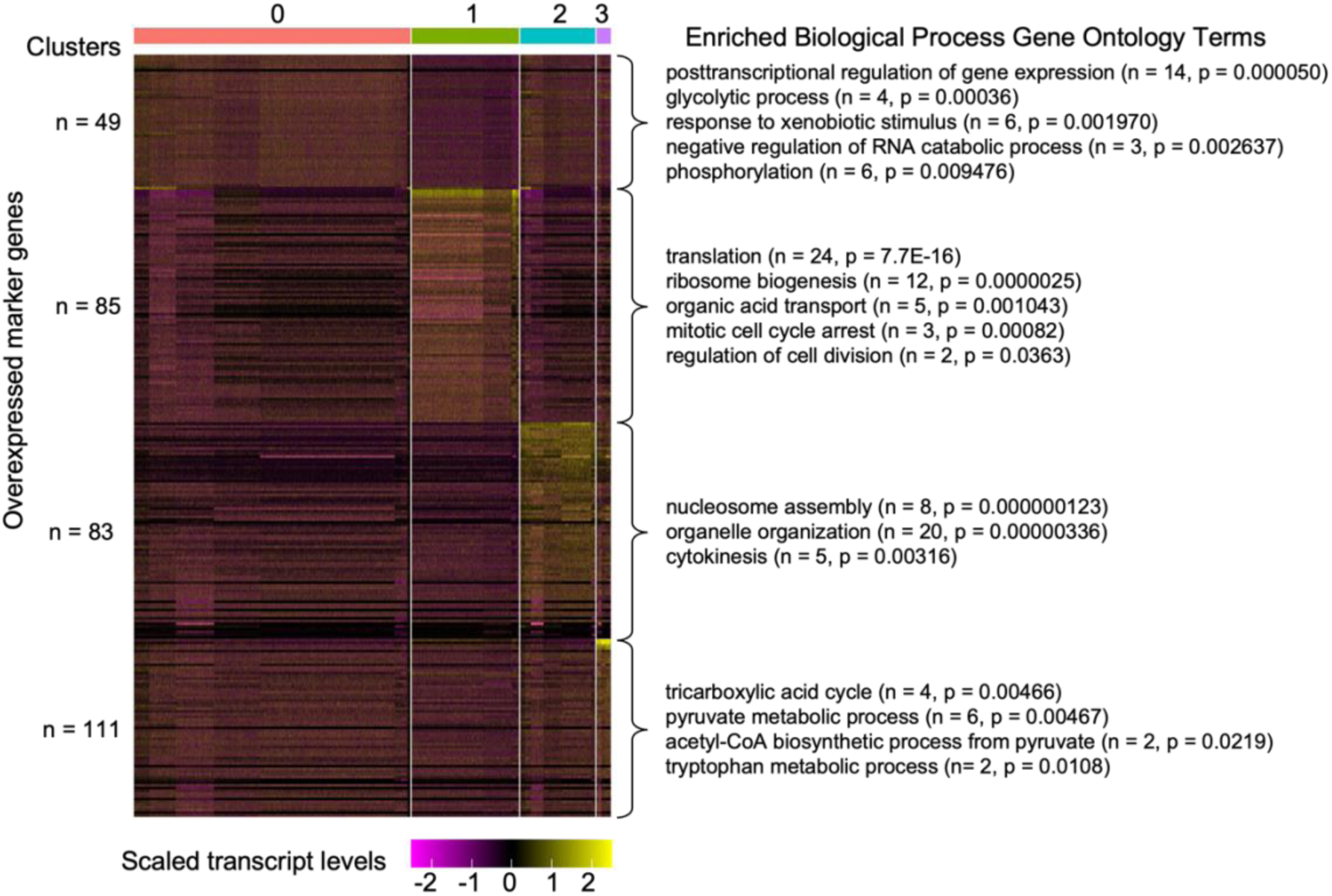
Gene markers of parasite clusters and associated GO terms. Heatmap shows scaled transcript levels for each marker genes separated per cluster. The number of markers for each cluster is indicated on the y-axis and cluster on the x-axis. For each top selected GO terms enriched for each set of markers is shown with the number of genes identified and adjusted p value.

**Supplementary figure 4.**
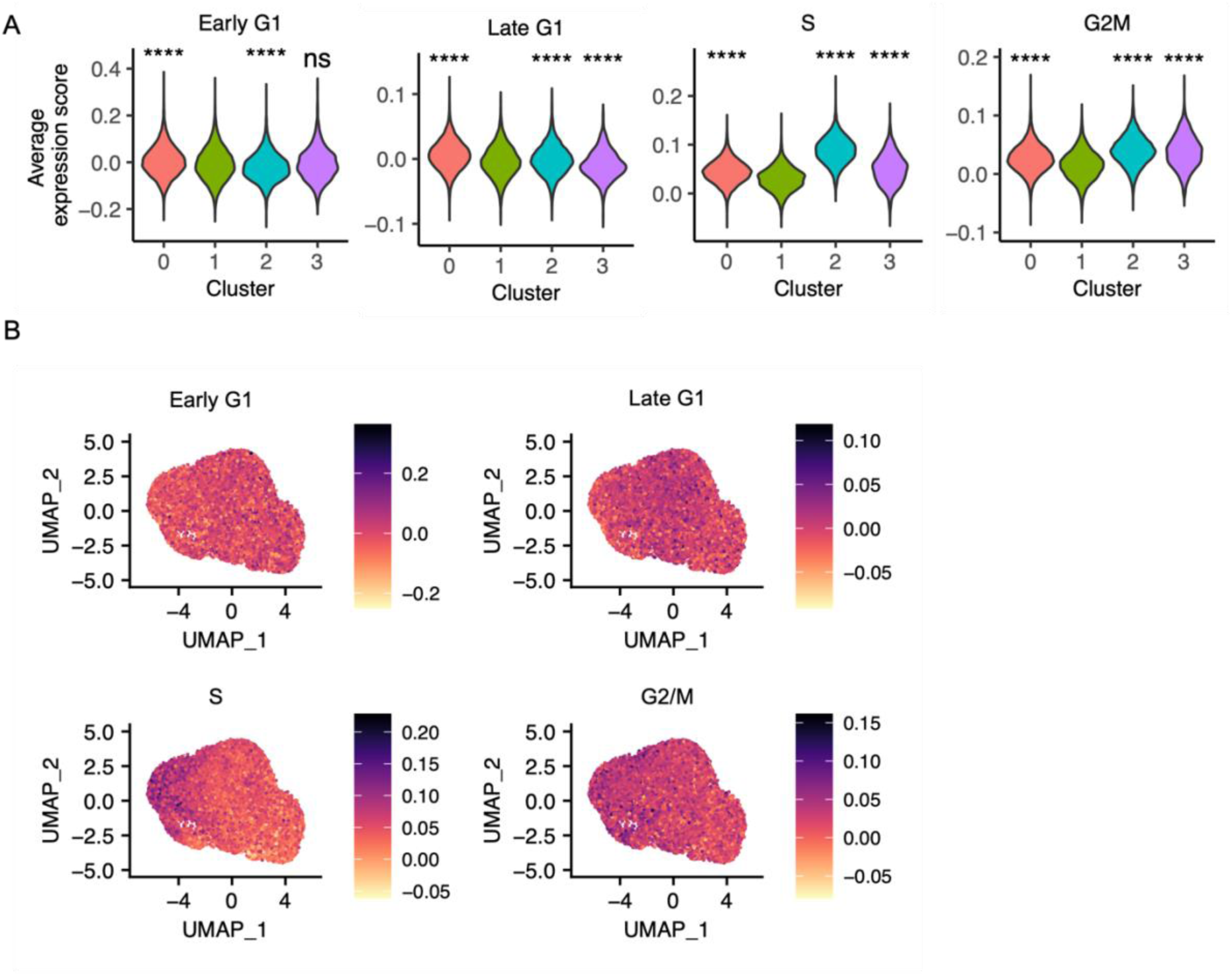
Cell cycle status of *T. brucei* in the cattle bloodstream inferred by transcriptome. a) Average gene expression score for Early G1, Late G1, S and G2/M phase marker genes. Individual t tests were performed using cluster 1 mean as the base, and compared to each other cluster. (ns: p > 0.05*: p <= 0.05 **: p <= 0.01 ***: p <= 0.001 ****: p <= 0.0001. b) UMAP of all cattle samples, coloured by each phase expression score.

**Supplementary figure 5.**
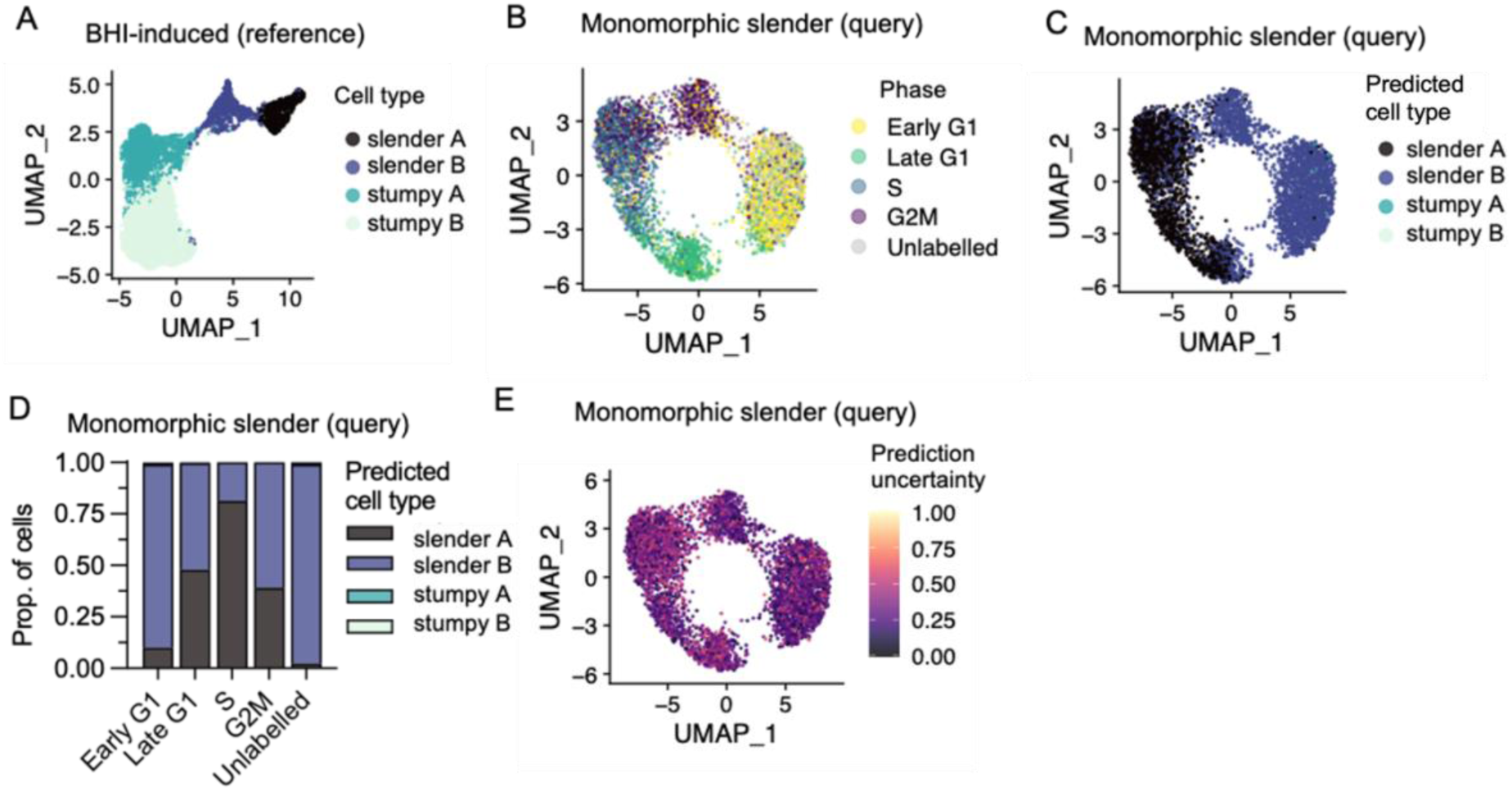
Cell type prediction correctly identifies slender parasites across datasets. **A**) UMAP of previously described scRNA-seq analysis of *T. brucei* treated with 10% brain heart infusion broth *in vitro* to generate slender and stumpy forms. B) UMAP of monomorphic slender forms *T. brucei* replicating in culture [24] used as query data, coloured by cell cycle phase. Monomorphs are unable to form arrested stumpy forms. C) UMAP of monomorphic slender forms coloured by predicted cell type. Labels and colours are consistent with reference data in a. D) Proportions of each monomorphic slender cell cycle phase (x-axis) as shown in 2b, predicted to be *in vitro* generated cell-types. E) UMAP of monomorphic slender form transcriptomes coloured by uncertainty in cell type predictions. 0 indicates low prediction uncertainty and 1 indicates high uncertainty.

**Supplementary figure 6.**
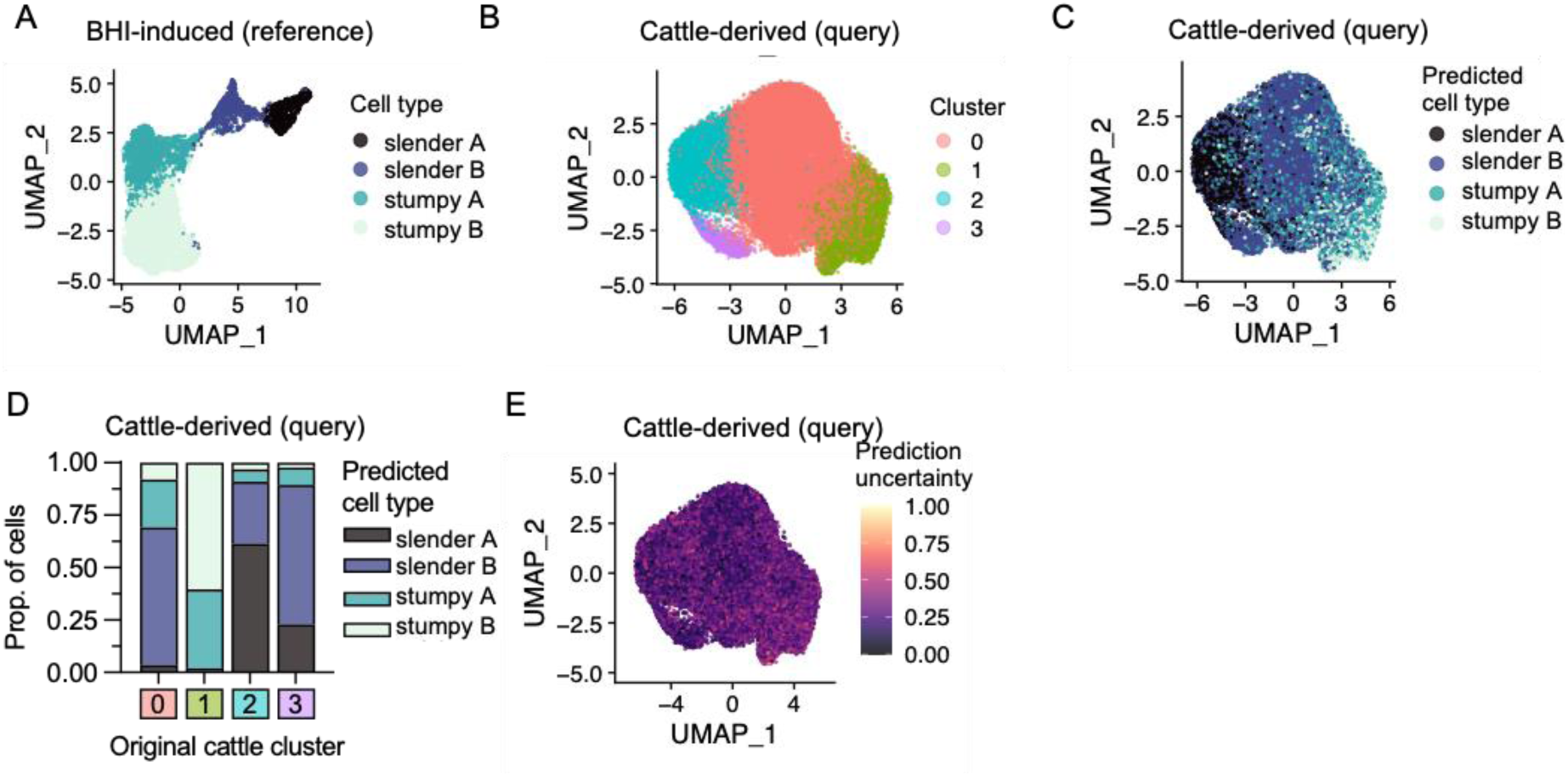
Cell type prediction indicates presenceof stumpy-like parasites in cattle bloodstream. A) UMAP of previously described scRNA-seq analysis of *T. brucei* treated with 10% brain heart infusion broth *in vitro* to generate slender and stumpy forms [15]. B) UMAP of all integrated cattle derived *T. brucei* used as query data for label transfer. Clusters are consistent with figure 1b. C) UMAP of cattle derived data coloured by predicted cell type. Labels and colours are consistent with reference data in 2a. D) Proportions of each cattle derived cluster (x-axis) as shown in 2b, predicted to be equivalent to *in vitro* generated cell-types. E) UMAP of all cattle derived transcriptomes coloured by uncertainty in cell type predictions. 0 indicates low prediction uncertainty and 1 indicates high uncertainty.

**Supplementary data 1.** Cluster markers (tab 1) and associated GO terms (tabs 2-5) for cattle data clusters 0-3 presented in Figure 1B.

**Supplementary data 2.** Results of differential expression analysis after pseudo-bulking of scRNA-seq data from mouse, cow and *in vitro* experiments presented in Figure 4.

Tab 1: Differential expression between mouse derived and BHI treated *in vitro T. brucei* samples.

Tab 2: Differential expression between cow derived and BHI treated *in vitro T. brucei* samples.

Tab 3: Differential expression between cow derived and mouse derived *T. brucei* samples.

Tab4: Differential expression of ESAG genes between cow and mouse derived samples.

Tab 5: Differential expression of ESAG4/GRESG4 genes.

Tab 6: GO term enrichment analysis of genes upregulated in cattle samples vs mouse samples.

Tab 7: GO term enrichment analysis of genes upregulated in mouse samples vs cattle samples.

**Supplementary data 3.** Fold change levels of slender and stumpy/stumpy-like marker gene transcripts in each dataset (cow, mouse and *in vitro*) presented in Figure 5.

## Notes

### Competing Interest Statement

The authors have declared no competing interest.

